# Crystal structure of SARS-CoV-2 nucleocapsid protein RNA binding domain reveals potential unique drug targeting sites

**DOI:** 10.1101/2020.03.06.977876

**Authors:** Sisi Kang, Mei Yang, Zhongsi Hong, Liping Zhang, Zhaoxia Huang, Xiaoxue Chen, Suhua He, Ziliang Zhou, Zhechong Zhou, Qiuyue Chen, Yan Yan, Changsheng Zhang, Hong Shan, Shoudeng Chen

## Abstract

The outbreak of coronavirus disease (COVID-19) in China caused by SARS-CoV-2 virus continually lead to worldwide human infections and deaths. It is currently no specific viral protein targeted therapeutics yet. Viral nucleocapsid protein is a potential antiviral drug target, serving multiple critical functions during the viral life cycle. However, the structural information of SARS-CoV-2 nucleocapsid protein is yet to be clear. Herein, we have determined the 2.7 Å crystal structure of the N-terminal RNA binding domain of SARS-CoV-2 nucleocapsid protein. Although overall structure is similar with other reported coronavirus nucleocapsid protein N-terminal domain, the surface electrostatic potential characteristics between them are distinct. Further comparison with mild virus type HCoV-OC43 equivalent domain demonstrates a unique potential RNA binding pocket alongside the β-sheet core. Complemented by *in vitro* binding studies, our data provide several atomic resolution features of SARS-CoV-2 nucleocapsid protein N-terminal domain, guiding the design of novel antiviral agents specific targeting to SARS-CoV-2.

## Introduction

The ongoing outbreak of coronavirus disease 2019 (COVID-19) is a new emerging human infectious disease caused by a novel coronavirus (<u>s</u>evere <u>a</u>cute <u>r</u>espiratory <u>s</u>yndrome <u>c</u>orona<u>v</u>irus <u>2</u>, SARS-CoV-2, previously known as 2019-nCoV). As of 28 February 2020, a cumulative total of 78,961 COVID-19 cases were confirmed with 2791 deaths in China. The emerged global epidemic spread rapidly with 4691 confirmed cases and 67 deaths across 51 countries outside of China (COVID-19 situation Report WHO, 28 Feb 2020). Despite remarkable efforts on containing spread of the virus, there is no specific targeted therapeutic currently.

SARS-CoV-2 is a betacoronavirus with single-stranded RNA genomes, like MERS-CoV and SARS-CoV. The first two-thirds of viral 30kb RNA genome, mainly named as ORF1a/b region, translates into two polyproteins (pp1a and pp1ab) and encodes most of the non-structural proteins (nsp). The rest parts of virus genome encode accessory proteins and four essential structural proteins, including spike (S) glycoprotein, small envelope (E) protein, matrix (M) protein, and nucleocapsid (N) protein^1,2^. Current antiviral drugs developed to treat coronavirus (CoV) infections primarily target S protein, the 3C-like (3CL) and papain-like (PLP) proteases^3,4^. Because mutant viruses in the S protein is prone to escape the targeted therapeutic with different host-cell receptor binding patterns^4^, as well as antibody-dependent enhancement (ADE) effects of S protein antibodies are found in MERS coronavirus^5^, there are several limitations on targeting S protein for antiviral approaches. Antiviral protease inhibitors may nonspecifically act on the cellular homologous protease, resulting in host cell toxicity and severe side effects. Therefore, novel antiviral strategies are needed to combat acute respiratory infections caused by this novel coronavirus SARS-CoV-2.

The CoV N protein is a multifunctional RNA-binding protein necessary for viral RNA transcription and replication. It plays many pivotal roles in forming helical ribonucleoproteins during packaging the RNA genome, regulating viral RNA synthesis during replication, transcription and modulating infected cell metabolism^6–8^. The primary functions of N protein are binding to the viral RNA genome, and packing them into a long helical nucleocapsid structure or ribonucleoprotein (RNP) complex^9,10^. *In vitro* and *in vivo* experiments revealed that N protein bound to leader RNA, and was critical for maintaining highly ordered RNA conformation suitable for replicating, and transcribing the viral genome^7,11^. More studies implicated that N protein regulated host-pathogen interactions, such as actin reorganization, host cell cycle progression, and apoptosis^12–14^. The N protein is also a highly immunogenic and abundantly expressed protein during infection, capable of inducing protective immune responses against SARS-CoV and SARS-CoV-2^15–18^.

The common domain architectures of coronavirus N protein are consisting of three distinct but highly conserved parts: An N-terminal RNA-binding domain (NTD), a C-terminal dimerization domain (CTD), and intrinsically disordered central Ser/Arg (SR)-rich linker. Previous studies have revealed that the NTD are responsible for RNA binding, CTD for oligomerization, and (SR)-rich linker for primary phosphorylation, respectively^19–23^. The crystal structures of SARS-CoV N-NTD^24^, infectious bronchitis virus (IBV) N-NTD^25,26^, HCoV-OC43 N-NTD^20^ and mouse hepatitis virus (MHV) N-NTD^27^ have been solved. The CoVs N-NTD have been found to associate with the 3' end of the viral RNA genome, possibly through electrostatic interactions. Additionally, several critical residues have been identified for RNA binding and virus infectivity in the N-terminal domain of coronavirus N proteins^24,27–29^. However, the structural and mechanistic basis for newly emerged novel SARS-CoV-2 N protein remain largely unknown. Understanding these aspects should facilitate the discovery of agents that specifically block the coronavirus replication, transcription and viral assembly^30^.

At present work, we report the crystal structure of SARS-CoV-2 nucleocapsid N-terminal domain (termed as SARS-CoV-2 N-NTD), as a model for understanding the molecular interactions that govern SARS-CoV-2 N-NTD binding to ribonucleotides. Compared with other solved CoVs N-NTD, we characterized the specificity surface electrostatic potential features of SARS-CoV-2 N-NTD. Additionally, we further demonstrated the unique RNA binding site characteristics. Our findings will aid in the development of new drugs that interfere with viral N protein and viral replication in SARS-CoV-2, and highly related virus SARS-CoV.

## Materials and methods

### Cloning, expression and purification

The SARS-CoV-2 N-FL plasmid is a gift from Guangdong Medical Laboratory Animal Center. We designed several constructs including: SARS-CoV-2 N-FL (residues from 1 to 419), SARS-CoV-2 N-NTD domain (residues from 41 to 174) and SARS-CoV-2 N-NTD domain (residues from 33 to 180) depending on secondary structure predictions and sequence conservation characteristics. The constructs were cloned into the pRSF-Duet-1 vector with N-terminal 6xHis-SUMO tag and expressed in E. coli strain *Rosetta*. SARS-CoV-2 N-NTD was induced with 0.1mM IPTG and incubated overnight at 16 °C in TB media. After Ni column chromatography followed by Ulp1 protease digestion for tag removal, SARS-CoV-2 N-NTD (41-174) proteins were further purified via size-exclusion chromatography (with buffer consisting of 20 mM Tris-HCl (pH 8.0), 150 mM sodium chloride, 1 mM dithiothreitol), and then concentrated by ultrafiltration to a final concentration of 22 mg/mL.

### Crystallization and data collection

Crystals were grown by the sitting drop method with 0.3ul protein (22 mg/mL) mixed with 0.6 μL and 0.3 μL well solution using Mosquito crystallization robot and after 3 days initial crystallization was performed under 16 °C under 20 mM sodium acetate, 100 mM sodium cacodylate (pH 6.5), 26 % PEG8000 conditions. Crystals were frozen in liquid nitrogen in reservoir solution supplemented with 15 % glycerol (v/v) as a cryoprotectant. X-ray diffraction data were collected at the South China Sea Institute of Oceanology, Chinese Academy of Sciences by Rigaku X-ray diffraction (XRD) instruments. The structure was solved by molecular replacement using PHENIX software suite^31^. The X-ray diffraction and structure refinement statistics are summarized in Table 1.

**Table 1.** Data collection and refinement statistics. Statistics for the highest-resolution shell are shown in parentheses.

|  | **SARS-CoV-2 N-NTD** |
| --- | --- |
| **Protein Data Bank code** | 6M3M |
| **Wavelength (Å)** | 1.5418 |
| **Resolution range** | 20.92 - 2.7 (2.796 - 2.7) |
| **Space group** | *P* 2<sub>1</sub> 2<sub>1</sub> 2<sub>1</sub> |
| **Unit cell**<br>(a, b, c, α, β, γ) | 58.88, 92.68, 97.32, 90, 90, 90 |
| **Total reflections** | 98913 (10077) |
| **Unique reflections** | 15133 (1481) |
| **Multiplicity** | 6.5 (6.8) |
| **Completeness (%)** | 99.55 (99.80) |
| **Mean I/sigma(I)** | 14.97 (2.9) |
| **Wilson B-factor** | 33.94 |
| **R-merge** | 0.1043 (0.3172) |
| **R-meas** | 0.1135 (0.3436) |
| **R-pim** | 0.04388 (0.1303) |
| **CC1/2** | 0.991 (0.96) |
| **CC\*** | 0.998 (0.99) |
| **Reflections used in refinement** | 15126 (1481) |
| **Reflections used for R-free** | 1514 (138) |
| **R-work** | 0.2578 (0.3551) |
| **R-free** | 0.2934 (0.4058) |
| **CC(work)** | 0.908 (0.692) |
| **CC(free)** | 0.851 (0.635) |
| **Number of non-hydrogen atoms** | 3952 |
| macromolecules | 3822 |
| solvent | 130 |
| **Protein residues** | 499 |
| **RMS(bonds)** | 0.004 |
| **RMS(angles)** | 0.72 |
| **Ramachandran favored (%)** | 96.48 |
| **Ramachandran allowed (%)** | 3.52 |
| **Ramachandran outliers (%)** | 0.00 |
| **Rotamer outliers (%)** | 0.00 |
| **Clashscore** | 11.15 |
| **Average B-factor** | 31.60 |
| macromolecules | 31.86 |
| solvent | 24.03 |

### SPR Analysis

Surface plasmon resonance (SPR) analysis was performed using a Biacore T200 with the CM5 sensor chip (GE Healthcare) at room temperature (25 °C). SARS-CoV-2 N-NTD (41–174) were exchanged into PBS buffer via gel-filtration. The CM5 chip surface was activated for 10 min using 0.2 M EDC/ 0.05 M NHS. After the injection of 30 μg/mL protein in 10 mM sodium acetate (pH 5.5) for three times to immobilize on one of channels of the CM5 chip up to ~5,800 response units, 10 μL of 1 M ethanolamine (pH 8.0) was used for blocking the remaining activated groups. Each of the analytes (AMP, GMP, UMP, CMP) were dissolve in PBS (pH 7.4, 0.05 % NP-20) and flowed through the chip surface at flow rate of 30 μL/min at 25 °C. 30 μL analytes were injected for affinity analysis with 60 s dissociation time. To understanding dose dependent affinity of analytes and SARS-CoV-2 N-NTD, we tested nine dilutions of analytes from 0.15625 mM to 10 mM. A blank flow channel was used as a reference channel to correct the bulk refractive index by subtracting the response signal of the reference channel from the signals of protein immobilized cells. The equilibrium constant (*K*_*D*_) for analytes binding to SARS-CoV-2 N-NTD was determined from the association and dissociation curves of the sensorgrams, using the BIAevaluation program (Biacore).

## Results

### Sequence features of SARS-CoV-2 nucleocapsid protein

It is reported that the complete genome of SARS-CoV-2 (MN908947, Wuhan-Hu-1 coronavirus) is 29.9 kb in length, similarly to 27.9 kb SARS-CoV and 30.1 kb MERS-CoV genome^32,33^ (Fig. 1A). Nucleocapsid (N) protein is translated from the 3’ end structural ORF^34–36^. According to Virus Variation Resource in National Center for Biotechnology Information databank^37^, SARS-CoV-2 N protein encoding region are conserved among the known NCBI 103 genome datasets. Only a few variations (S194L in virus strain Foshan/20SF207/2020, K249I in virus strain Wuhan/IVDC-HB-envF13-21/2020, P344S in virus strain Guangzhou/20SF206/2020) in N protein are found in public genomic epidemiology. An overall domain architecture of N protein among four coronaviruses (SARS-CoV-2, SARS-CoV, MERS-CoV and HCoV-OC43) are shown in Figure 1B, which indicates that SARS-CoV-2 shares typical characteristics with other coronaviruses. Zoom into the completed genomic sequence of SARS-CoV-2 N protein encoding region, we find that the sequence identities between SARS-CoV-2 with SARS-CoV, MERS-CoV, and HCoV-OC43 are 89.74%, 48.59%, 35.62%, respectively (Fig. S1)^38,39^. Since full-length SARS-CoV-2 N protein aggregated status were found in our expression and purification studies (Fig. S2), as well as previous reported data on other coronavirus nucleocapsid protein, we next investigate the structural studies on N-terminal region of SARS-CoV-2 N protein (termed as SARS-CoV-2 N-NTD).

**Figure 1:**
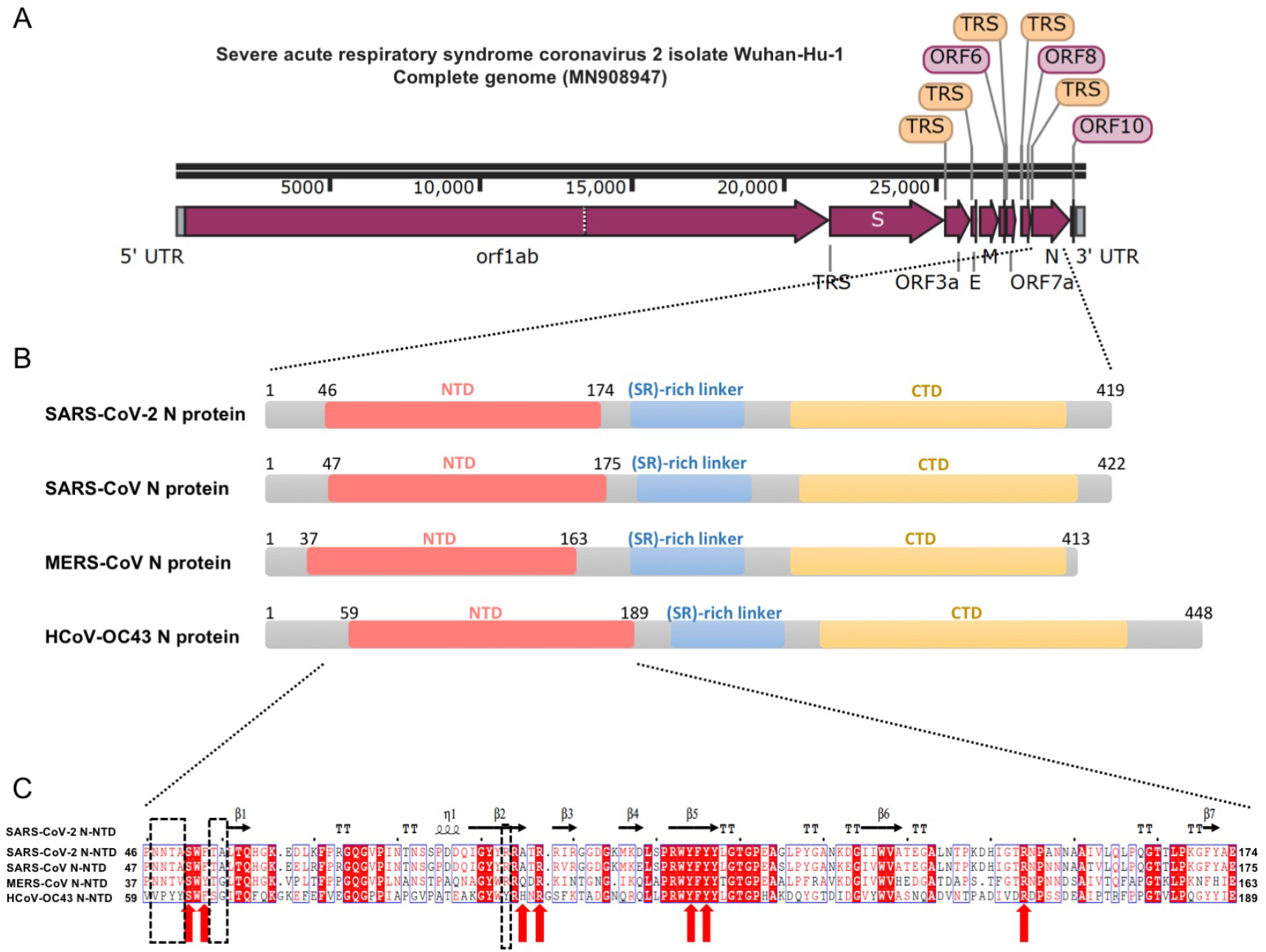
Sequence features of SARS-CoV-2 nucleocapsid protein. **Figure legend**: **A.** The complete whole genomic features of SARS-CoV-2 isolate Wuhan-Hu-1 (Genebank: MN908947). UTR: untranslated region; orf/ORF: open reading frame; TRS: transcriptional regulatory sequences; S: spike glycoprotein encoding region; E: envelope protein encoding region; M: membrane protein encoding region; N: nucleocapsid protein encoding region. The figure is illustrated by SnapGene Viewer; **B.** Domain architectures of coronavirus nucleocapsid protein. NTD: N-terminal RNA-binding domain; CTD: C-terminal dimerization domain; **C.** Multiple sequence alignment of SARS-CoV-2 N-NTD with SARS-CoV N-NTD (UniProtKB: P59595), MERS-CoV N-NTD (UniProtKB: R9UM87), HCoV-OC43 N-NTD (UniProtKB: P33469). Red arrows indicate conserved residues for ribonucleotide binding site, dash boxes indicate variably residues in the structural comparisons.

### Crystal structure of SARS-CoV-2 N-NTD

In order to obtain the atomic information of SARS-CoV-2 N-NTD, we solve the structure at 2.7 Å resolution using X-ray crystallography technology. Briefly, 47-173 residues of SARS-CoV-2 N protein were constructed, expressed and purified as described protocol (Materials and Methods). The structure of SARS-CoV-2 N-NTD was determined by molecular replacement using the SARS-CoV N-NTD structure (PDB:2OG3) as the search model^24^. The final structure was refined to R-factor and R-free values of 0.26 and 0.29, respectively. The complete statistics for data collection, phasing and refinement are presented in Table 1. Unlike to SARS-CoV N-NTD crystals packing modes (monoclinic form at 2OFZ, cubic form at 2OG3)^24^, SARS-CoV-2 N-NTD crystal shows orthorhombic crystal packing form with four N-NTD monomers in one asymmetry unit (Fig. 2A). Although lacks of evidence for real RNP organization in the mature virions, the differences in the crystal packing patterns may implicate other potential contacts in SARS-CoV-2 RNP formation process. All four monomers in one asymmetric unit of the SARS-CoV-2 N-NTD crystal structure shared similar right-handed (loops)-(β-sheet core)-(loops) sandwiched structure, as conserved among the CoVs N-NTD (Fig. 2B). The β-sheet core is consisted of five antiparallel β-strands with a single short 3_10_ helix just before strand β2, and a protruding β-hairpin between strands β2 and β5. The β-hairpin is functionally important for CoV N-NTD, implicated in mutational analysis of amino acid residues for RNA binding^29^ (Fig. 2C). The SARS-CoV-2 N-NTD is enriched in aromatic and basic residues, folding into a hand shape resembles with basic fingers that extend far beyond the β-sheet core, a basic palm, and an acidic wrist (Fig. 2D).

**Figure 2:**
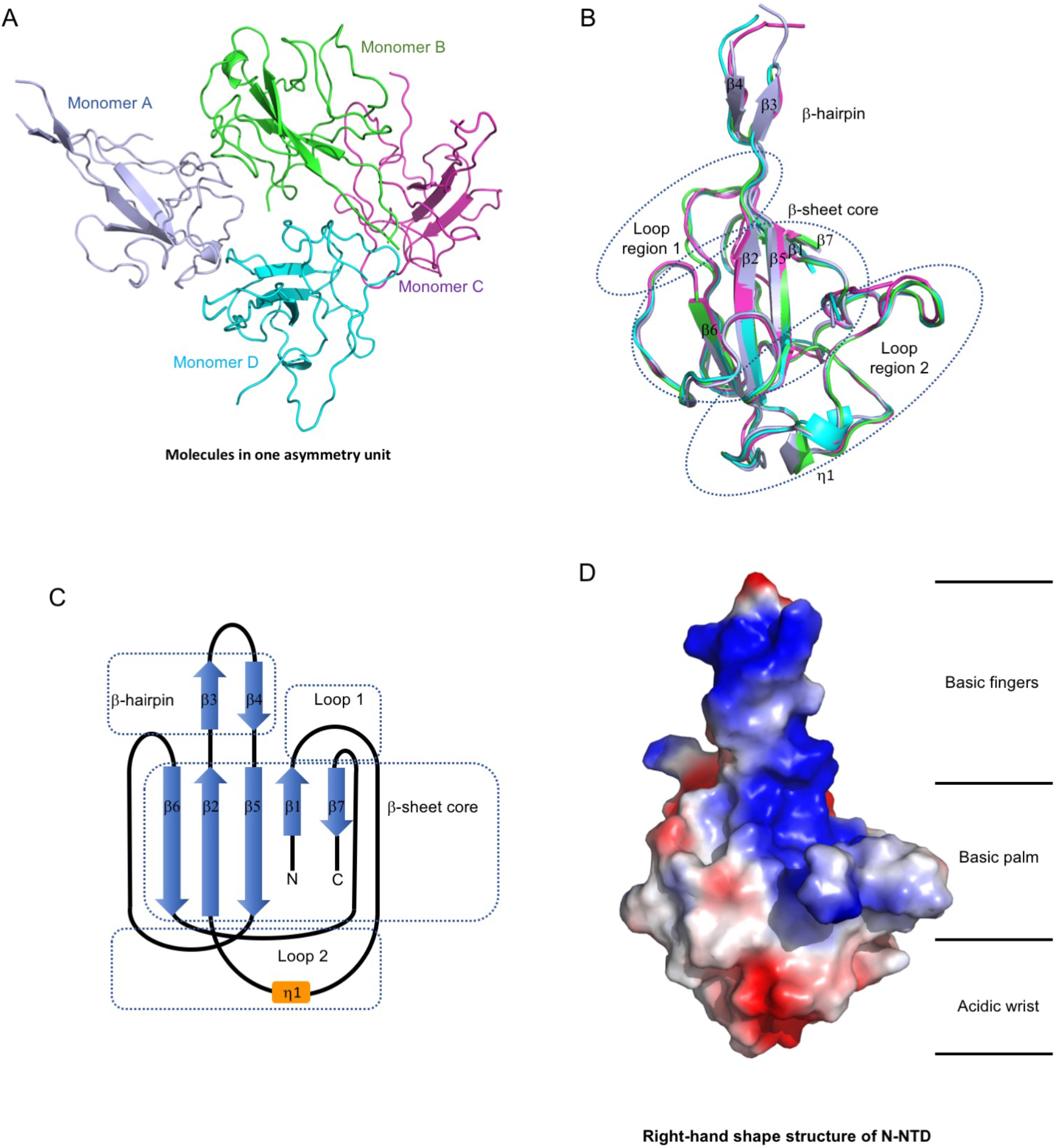
Structural overview of SARS-CoV-2 N-NTD. **Figure legend**: **A.** Ribbon representation of SARS-CoV-2 N-NTD molecules in one asymmetric unit. The four molecules are highlighted with different color respectively; **B.** Superimpositions of four molecules in one asymmetric unit. The dash circles indicate the sandwiched structure composed of Loop region 1, β-sheet core, and Loop region 2. The β-strand is labeled with β1 to β7, the 3_10_ helix is labeled with η1; **C.** Topological style illustration of SARS-CoV-2 N-NTD structure; **D.** Electrostatic surface of the SARS-CoV-2 N-NTD. Blue denotes positive charge potential, while red indicates negative charge potential. The potential distribution was calculated by Pymol. The values range from −5 kT (red) to 0 (white) and to +5 kT, where k is the Boltzmann constant and T is the temperature.

### Comparison of SARS-CoV-2 N-NTD with related viral N-NTD structures

To obtain more specific information, we first mapped the conserved residues between SARS-CoV-2 N-NTD with SARS-CoV N-NTD, MERS-CoV N-NTD, HCoV-OC443 N-NTD, respectively (Fig 3A). The most conserved residues distribute on the basic palm region (Fig. 3A, blue and green region), while the less conserved residues locate in basic fingers and acidic wrist region (Fig. 3A, pink and red region). The available CoVs N-NTD crystal structures allowed us to compare the electrostatic potential on the surface. As shown in Fig. 3B, although CoV N-NTDs all adapted similar overall organizations, the surface charge distribution patterns are different. Consisting with our observations, previous modeling of related coronaviral N-NTDs also shown markedly differ in surface charge distributions^24^. Superimposition of SARS-CoV-2 N-NTD with three kinds of CoVs N-NTD are shown in Fig. 3C. Compared with SARS-CoV N-NTD, SARS-CoV-2 N-NTD show a 2.1 Å movement in the β-hairpin region forward to nucleotide binding site (Fig. 3C, left panel). While compared with MERS-CoV N-NTD, SARS-CoV-2 N-NTD show a less extended β-hairpin region, and a distinct relax N-treminal tail (Fig. 3C, middle panel). In consistently, SARS-CoV-2 N-NTD show a distinct relax N-treminal tail, and a 2 Å movement in the β-hairpin region backward to the opposite side of nucleotide binding site when the structure is compared with HCoV-OC43 N-NTD (Fig. 3C, right panel). These differences dramatically change the surface characterizations of the protein, may result in the RNA-binding cleft being adaptive to its own RNA genome.

**Figure 3:**
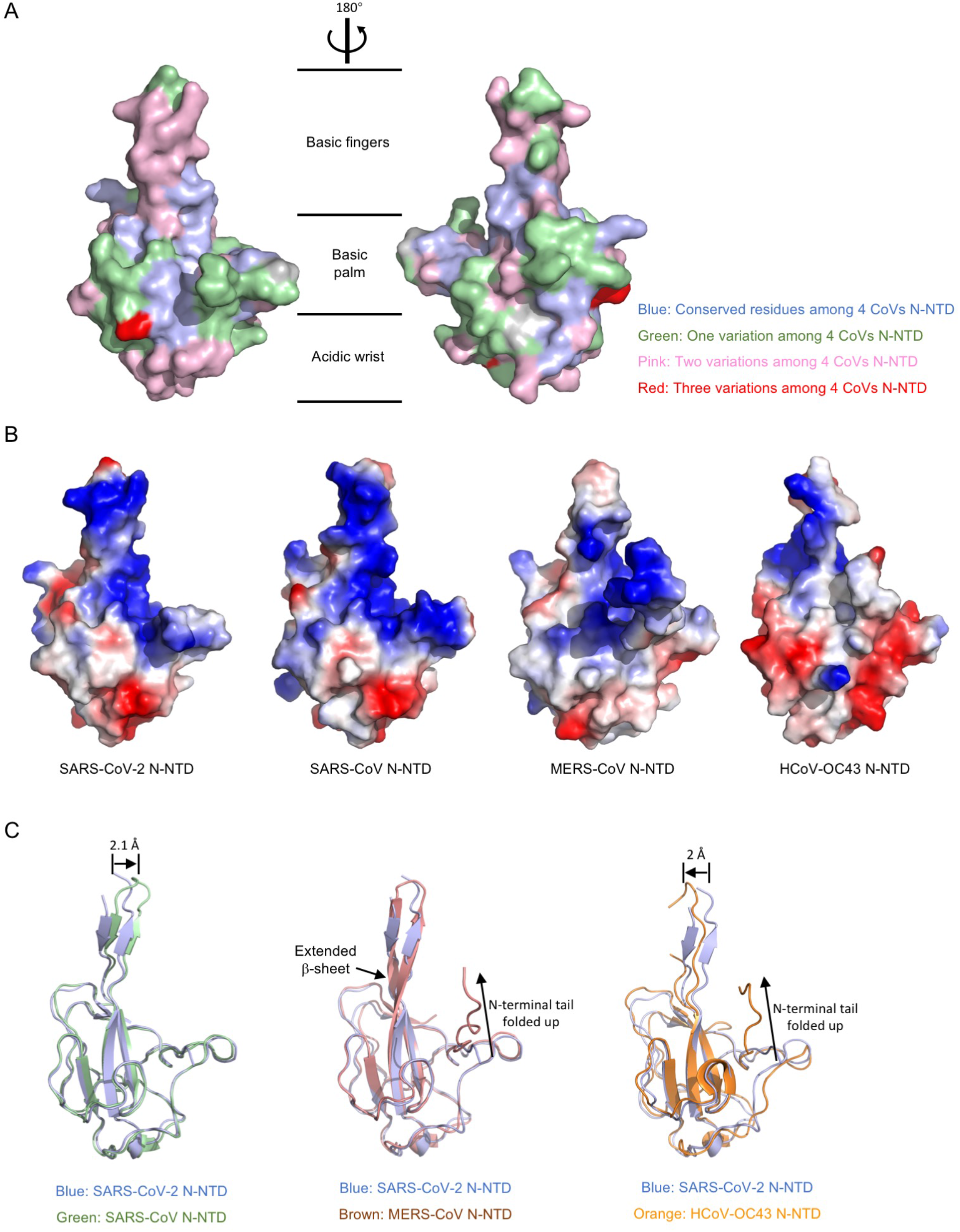
Comparison of SARS-CoV-2 N-NTD with related viral N-NTD structures. **Figure legend**: **A.** Mapping the conserved surfaces of four CoV N-NTDs in SARS-CoV-2 N-NTD structure. The multiple sequence alignment used for mapping is shown in Fig. 1C. Blue denotes conserved residues among 4 CoVs N-NTD; green denotes one variation among 4 CoVs N-NTD; pink denotes two variations among 4 CoVs N-NTD; red denotes three variations among 4 CoVs N-NTD; **B.** Electrostatic surface of the SARS-CoV-2 N-NTD, SARS-CoV N-NTD, MERS-CoV N-NTD, HCoV-OC43 N-NTD. Blue denotes positive charge potential, while red indicates negative charge potential; **C.** Overall structural comparison of SARS-CoV-2 N-NTD with related viral N-NTD structures. Left: superimposition of SARS-CoV-2 N-NTD (blue) to SARS-CoV N-NTD (green); middle: superimposition of SARS-CoV-2 N-NTD (blue) to MERS-CoV N-NTD (brown); right: superimposition of SARS-CoV-2 N-NTD (blue) to HCoV-OC43 N-NTD (orange).

### A potential unique drug target pocket in SARS-CoV-2 N-NTD

Although there are several CoV N-NTDs structures have been solved, the structural basis for ribonucleoside 5’-monophosphate binding of N protein had only been described in HCoV-OC43, a relative type typically causing mild cold symptoms^40^. Since the surface characterizations of N-NTD between SARS-CoV-2 with HCoV-OC43 are distinct, we next explored the differences of RNA binding mechanistic basis with superimposition analysis. Previous studies had shown that, HCoV-OC43 N-NTD contained Adenosine monophosphate (AMP)/ uridine monophosphate (UMP)/ cytosine monophosphate (CMP)/ guanosine monophosphate (GMP) binding site alongside the middle two β strands of its β-sheets core^40^. In the complex structure of HCoV-OC43 N-NTD with ribonucleotides, the phosphate group was bound by Arg 122 and Gly 68 via ionic interactions, the pentose sugar ribose 2’-hydroxyl group was recognized by Ser 64 and Arg164, the nitrogenous base inserted into a hydrophobic pocket consisting of Phe 57, Pro61, Tyr 63, Tyr102, Tyr 124, and Tyr 126, mainly interacted with Tyr 124 via π-π stacking forces (Fig. S3). It is proposed that this ribonucleotide binding mechanism are essential for all coronavirus N proteins, applying to develop CoV N-NTD-target agents.

To obtain the structure information of SARS-CoV-2 N-NTD ribonucleotide binding site, we make a superimposition of SARS-CoV-2 N-NTD with HCoV-OC43 N-NTD-AMP complex. As expectedly, the root mean square deviation (RMSD) between these two structure coordinates are 1.4 Å over 136 superimposed Cα atoms. However, a number of difference around the ribonucleotide binding site were shown as superimposition of SARS-CoV-2 N-NTD with HCoV-OC43 N-NTD. The major difference is N-terminal tail of N-NTD with sequence variation (SARS-CoV-2: 48 <u>**NNTA**</u> 51 versus HCoV-OC43: 60 <u>**VPYY**</u> 63). In HCoV-OC43 N-NTD, the tail folded up to compose a nitrogenous base binding channel, whereas this region extended outward in SARS-CoV-2 one (Fig. 4A). The N-terminal tail movement contributed to the change of N-NTD surface charge distribution, at which nucleotide binding cavity became easier to accessible in SARS-CoV-2 N-NTD (Fig. 4B and C). The second difference is on phosphate group binding site, which SARS-CoV-2 N-NTD have larger sidechain residues (55 <u>**TA**</u> 56) compared with HCoV-OC43 N-NTD equivalents (67 <u>**SG**</u> 68) (Fig. 4D). Structural superimposition suggested additional polar properties of Thr 55 and Ala 56 in SARS-CoV-2 N-NTD may increase the steric clash with ribonucleotide phosphate moiety (Fig. 4E and F). The third difference is on the edge of nitrogenous base recognized hydrophobic pocket, where SARS-CoV-2 N-NTD had Arg 89 residues compared with HCoV-OC43 N-NTD Tyr 102 equivalents (Fig. 4G). The change of these residues sidechain may lead to dramatic decreasing of non-polar properties and increasing of polar properties in the nitrogenous base binding site (Fig. 4H and I). To evaluate these different observations in our structure, Surface plasmon resonance (SPR) analysis experiments were next performed to assess the binding affinity between SARS-CoV-2 N-NTD with all four kinds of ribonucleotide AMP/UMP/CMP/GMP. Intriguingly, all ribonucleoside 5’-monophosphate, excepted for GMP (*K*_*D*_ value is 8 mM), shown little binding signals in assays (Fig. S4). Taken together, the above results suggested a potential distinct RNA binding patterns between SARS-CoV-2 N protein with HCoV-OC43 N protein.

**Figure 4:**
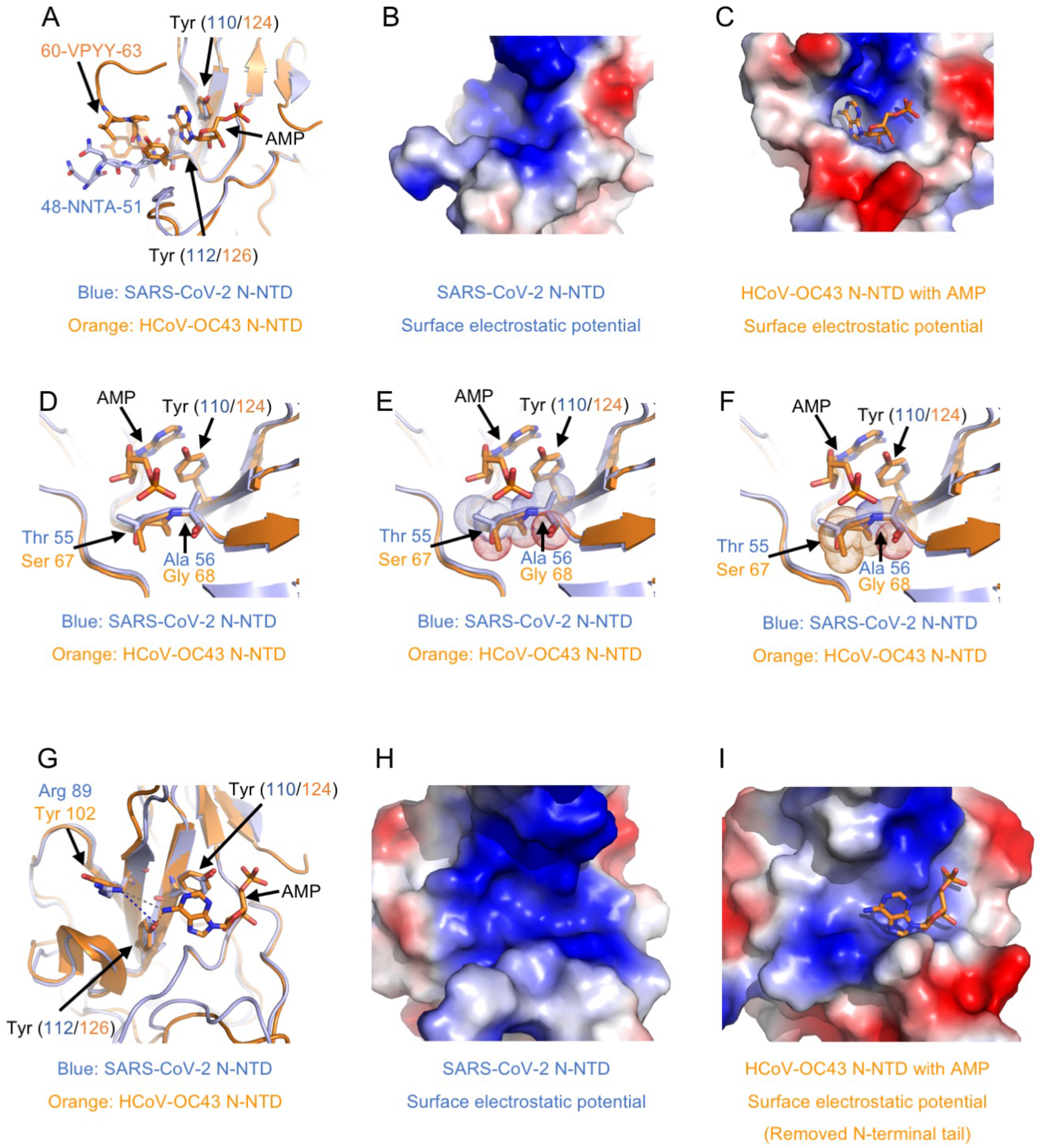
A potential unique drug target pocket in SARS-CoV-2 N-NTD. **Figure legend**: **A.** Detailed view of ribonucleotide binding pocket in superimposition structures between SARS-CoV-2 N-NTD with HCoV-OC43 N-NTD AMP complex. AMP, interacting residues and equivalents are highlighted with stick representation; **B.** Electrostatic surface of the potential ribonucleotide binding pocket on SARS-CoV-2 N-NTD; **C.** Electrostatic surface of the ribonucleotide binding pocket on HCoV-OC43 N-NTD; **D.** Detailed view of phosphate group binding site in superimposition structures between SARS-CoV-2 N-NTD with HCoV-OC43 N-NTD AMP complex; **E.** Dot representation of SARS-CoV-2 residues Thr 55 and Ala 56, which indicates potential steric clashes with the ribonucleotide phosphate group; **F.** Dot representation of HCoV-OC43 N-NTD residues Ser 67 and Gly 68; **G.** Detailed view of nitrogenous base binding site in superimposition structures between SARS-CoV-2 N-NTD with HCoV-OC43 N-NTD AMP complex; **H.** Electrostatic surface of the potential ribonucleotide nitrogenous base binding pocket on SARS-CoV-2 N-NTD; **I.** Electrostatic surface of the ribonucleotide nitrogenous base binding pocket on HCoV-OC43 N-NTD. In electrostatic surface potential panels, blue denotes positive charge potential, while red indicates negative charge potential. The potential distribution was calculated by Pymol. The values range from −5 kT (red) to 0 (white) and to +5 kT, where k is the Boltzmann constant and T is the temperature.

## Discussion

Structure-based drug discovery has been shown to be an advance approach for the development of new therapeutics. Many ongoing studies are developed to treat COVID-19 primarily targeting the spike protein, viral proteases (3C-like protease and papain-like protease). However, there is little effective targeted therapeutic currently. Recent studies demonstrated that N proteins will be a good drug-targeting candidate in other CoVs since they process several critical functions, such as RNA genomic packing, viral transcription and assembly, in the infectious cell^10^. However, the molecular basis of SARS-CoV-2 N protein is yet to be elucidated. Here, we present the 2.7 Å crystal structure of SARS-CoV-2 N protein N-terminal domain, revealing the specific surface charge distributions which may facilitate drug discovery specifically to SARS-CoV-2 N protein ribonucleotide binding domain.

On the structural basis of SARS-CoV-2 N-NTD, several residues in the ribonucleotide binding domain were found to distinctly recognize the CoV RNA substrates. The N-terminal tail residues (Asn 48, Asn 49, Thr 50, and Ala 51) is more flexible and extended outward compared with equivalent residues in HCoV-OC43 N-NTD, possibly opening up the binding pocket into fitting with viral RNA genomic high order structure. Residues Arg 89, instead of HCoV-OC43 N-NTD Tyr 102, may contribute to guanosine base recognition despite the overall ribonucleotide binding may be excluded by residues Thr 55 and Ala 56 in the phosphate moiety recognition site.

Up to date, seven coronaviruses have been identified as human-susceptible virus, among which HCoV-229E, HCoV-NL63, HCoV-HKU1 and HCoV-OC43 with low pathogenicity cause mild respiratory symptoms similar to common cold, whereas the other three betacoronaviruses, SARS-CoV-2, SARS-CoV and MERS-CoV lead to severe and potential fatal respiratory tract infections^32,41,42^. Previous study reported the structural basis of HCoV-OC43 N-NTD with AMP, GMP, UMP, CMP and a virtual screening-base compound PJ34. However, our data suggested that SARS-CoV-2 employed a unique pattern for binding RNA with atomic resolution information. The structure not only help us to understand the RNA-binding mechanisms between severe infectious coronavirus with mild infectious one, but also guide the design of novel antiviral agents specific targeting to SARS-CoV-2.

## Acknowledgments

We thank for Guangdong Medical Laboratory Animal Center for providing the N-protein encoding gene plasmids, Dr. Yongzhi Lu from Guangzhou Institutes of Biomedicine and Health (Chinese Academy of Sciences) for the initial crystals X-ray diffraction screening, supports from Dr. Xuan Ma of South China Sea Institute of Oceanology (Chinese Academy of Sciences) for home source X-ray diffraction facility. This work was supported by National Natural Science Foundation of China (31770801) and Special Fund for Scientific and Technological Innovation Strategy of Guangdong Province of China (2018B030306029, 2017A030313145) to S.C.

## Conflict of interest

The authors declare no conflict of interest

## Data Availability Statement

The structures in this paper are deposited to the Protein Data Bank with 6M3M access code.

## Supplementary Figures

**Figure S1:**
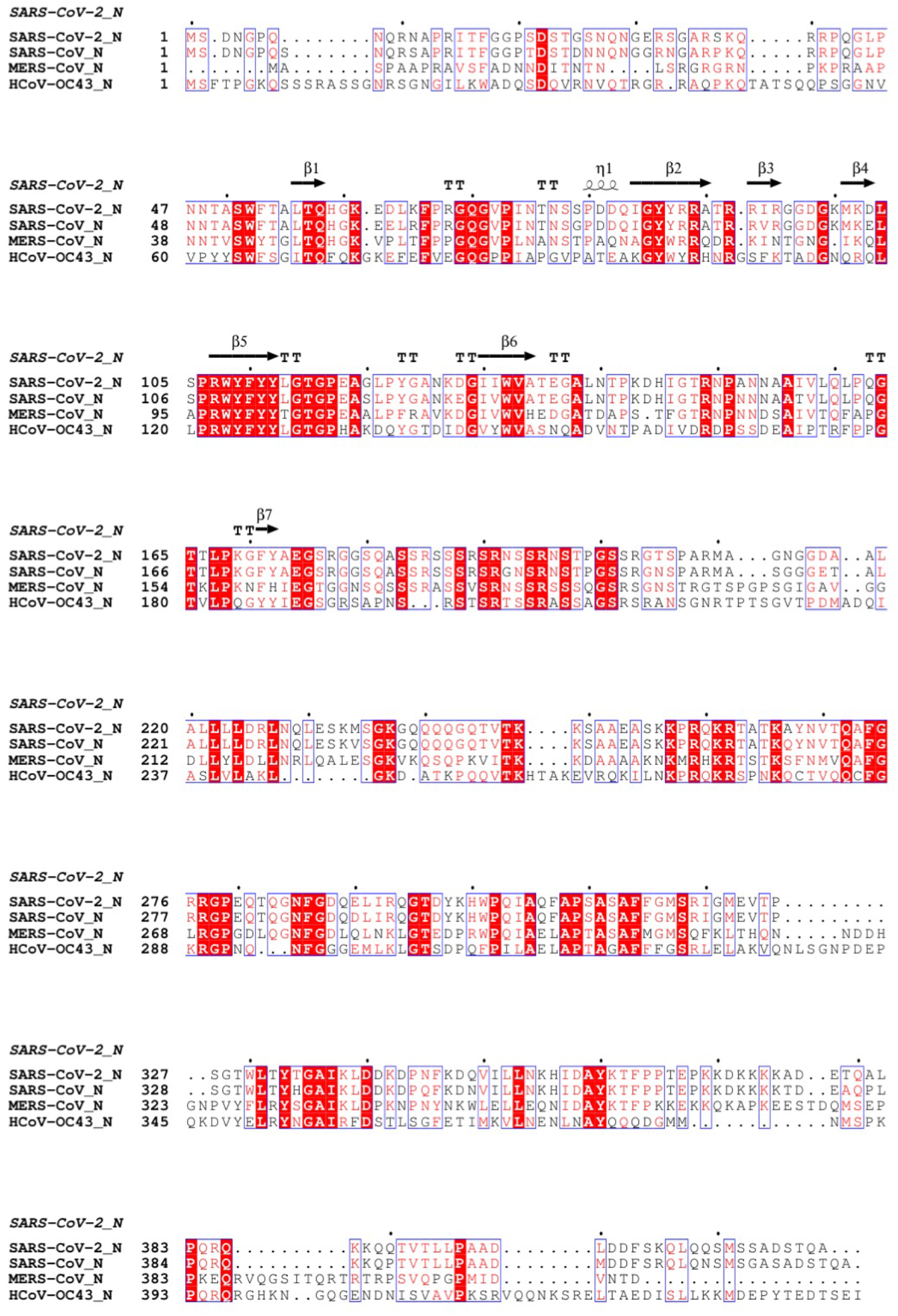
Multiple sequence alignment of coronavirus nucleocapsid protein. **Figure legend**: The alignment is accomplished at online server (Clustal Omega), and illustrated with ESPript 3.0 server. The top line with SARS-CoV-2_N label indicates secondary structure elements extracted from structural coordinate.

**Figure S2:**
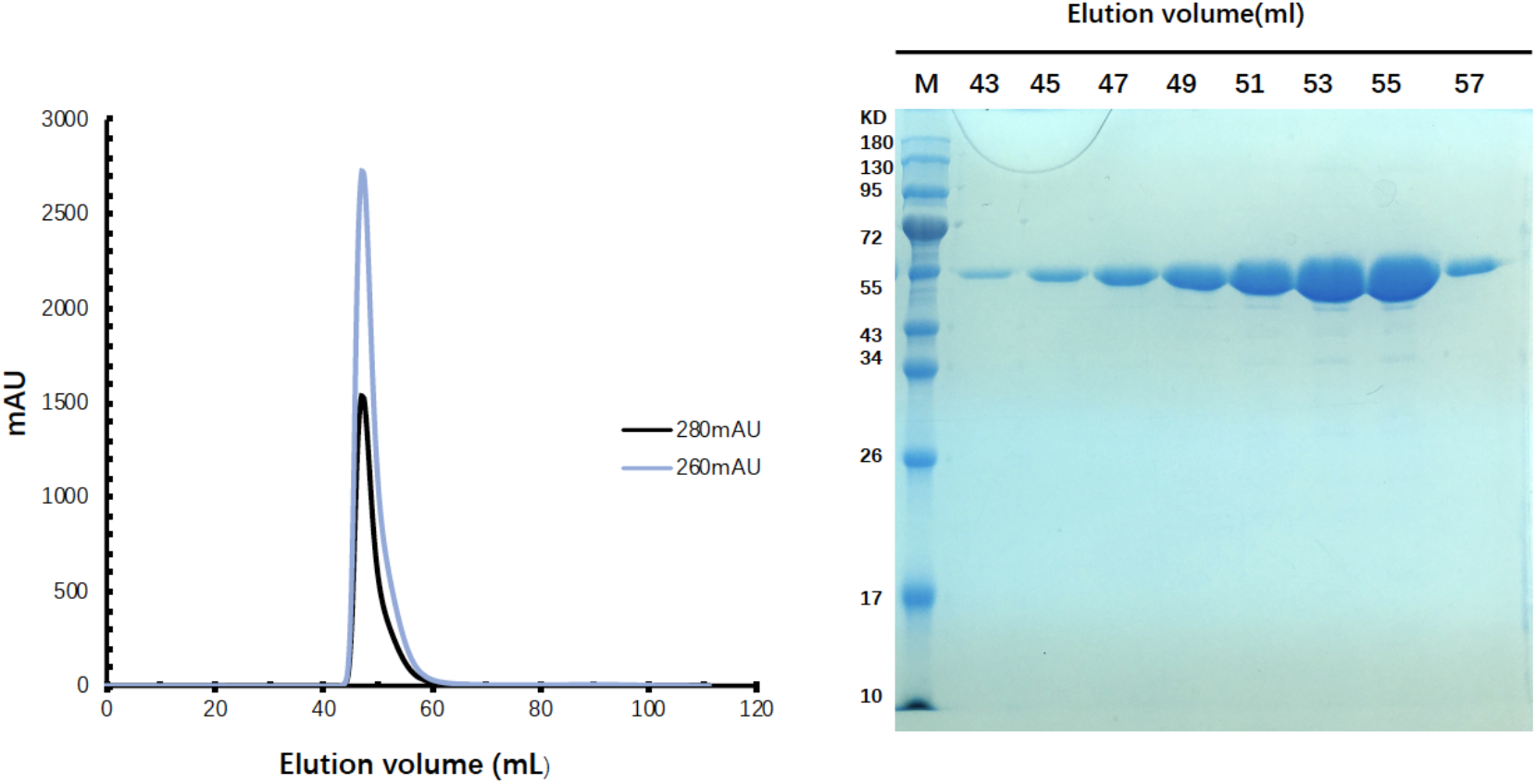
Full-length nucleocapsid protein purification. **Figure legend** : On the Left panel, a gel filtration chromatography result (HiLoad® Superdex® 200 pg GE Healthcare) of recombined full-length nucleocapsid protein. On the right panel, a 12% SDS-PAGE electrophoresis is used to analyze the purification samples.

**Figure S3:**
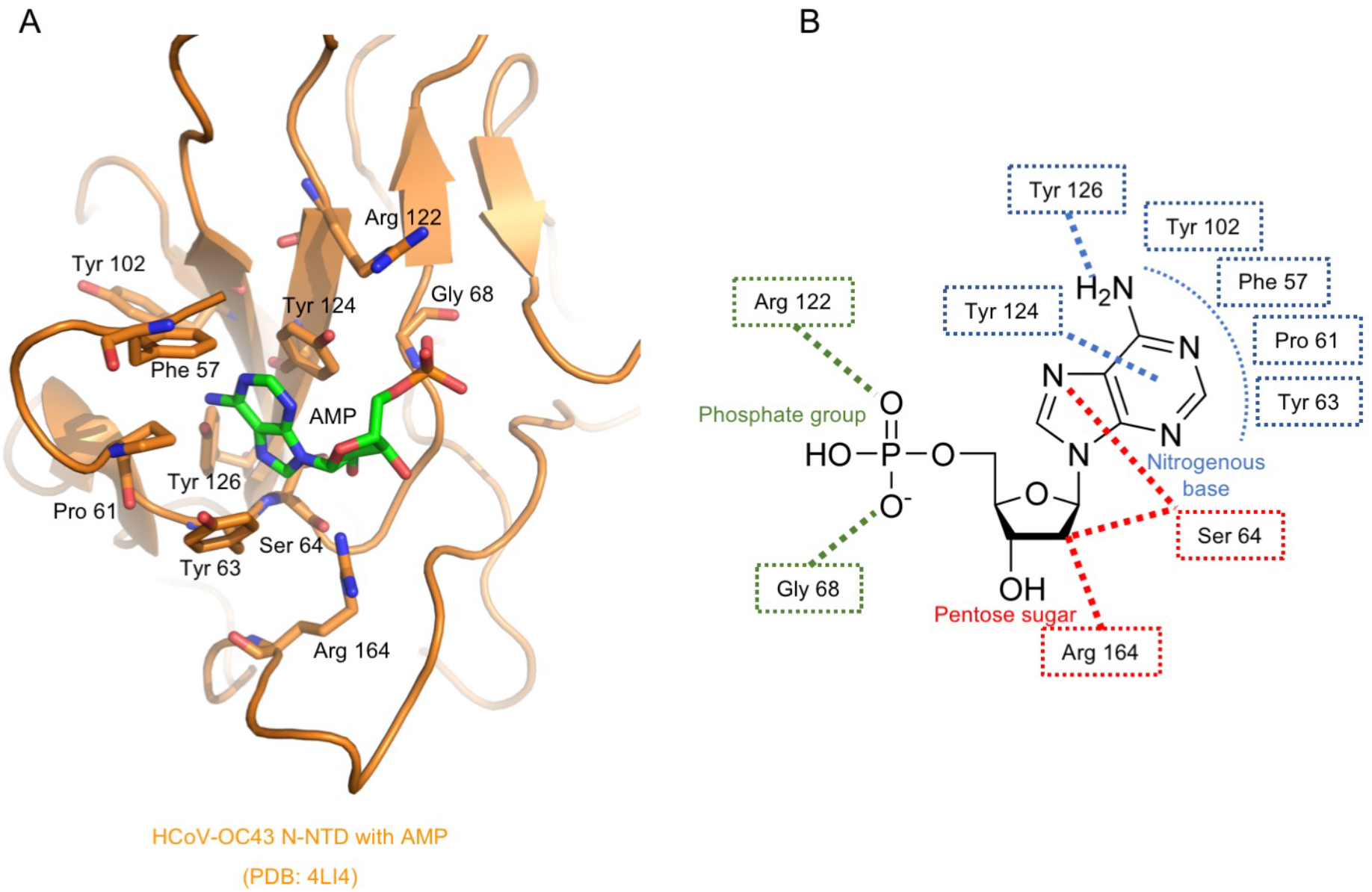
Ribonucleotide binding pocket in HCoV-OC43 N-NTD AMP complex. **Figure legend: A.** Detailed view of ribonucleotide binding pocket in HCoV-OC43 N-NTD AMP complex. AMP and its interacting residues are highlighted with stick representation; **B.** Simple illustration of AMP binding pocket. Nitrogenous base and its binding residues are color with blue. Pentose sugar and its binding residues are color with red. Phosphate group and its binding residues are color with green.

**Figure S4:**
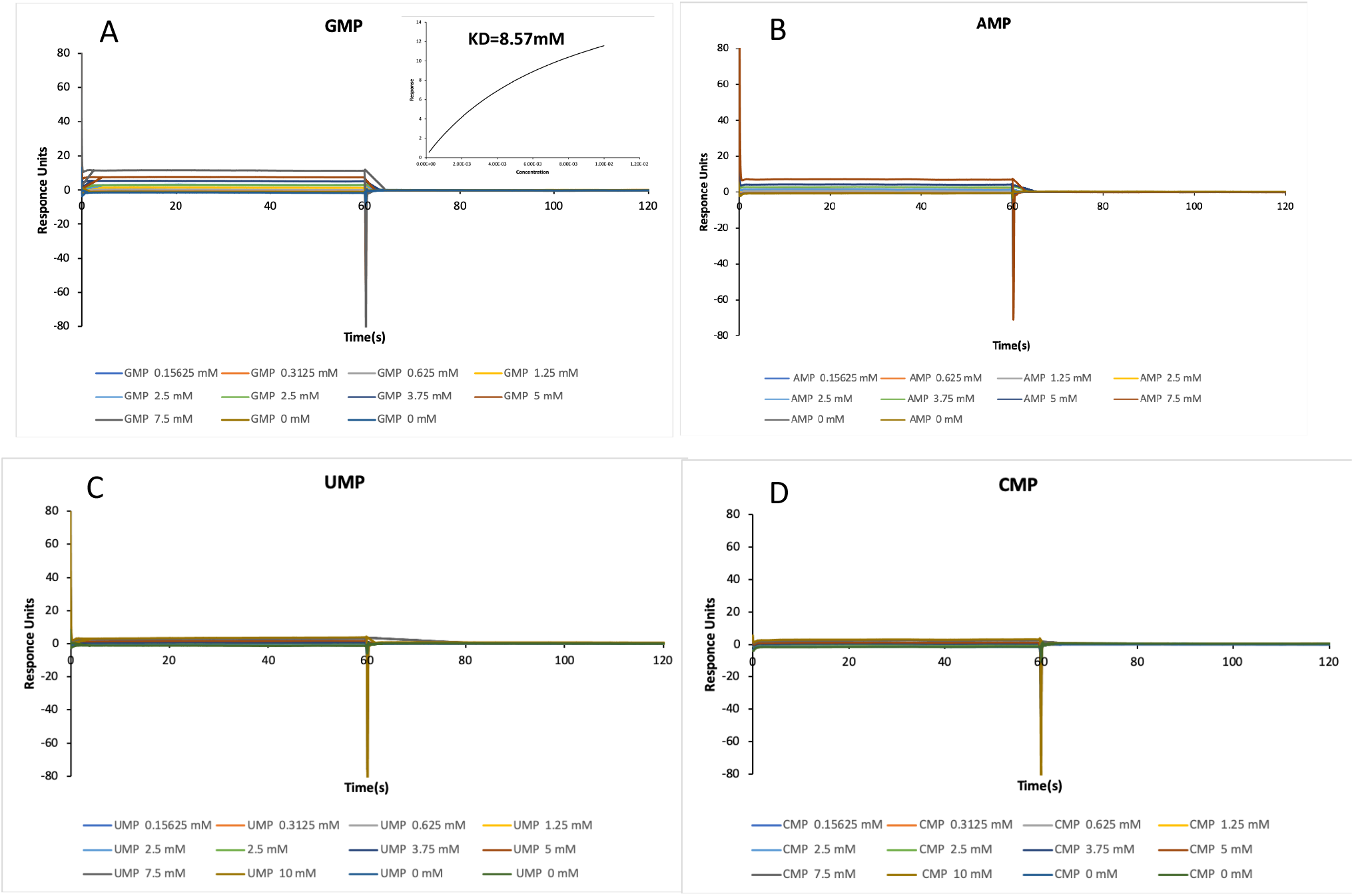
*In vitro* SPR assays for SARS-CoV-2 N-NTD with ribonucleotides. **Figure legend: A.** SPR sensorgram of the binding of varying concentrations of GMP (0, 0.15625, 0.3125, 0.625, 1.25, 2.5, 3.75, 5, 7.5 mM) to SARS-CoV-2 N-NTD captured CM5 chip. The curve in up-right box represents the binding affinity by fitting the sensorgram to a Langumir binding rate equations. The similar SPR sensorgrams of the binding of AMP, UMP, and CMP are shown in panel **B**, **C**, and **D**, respectively.

## Reference

1. Brian DA & Baric RS. Coronavirus genome structure and replication. Current Topics in Microbiology and Immunology 2005; 287 :1–30.

2. Cui J, Li F & Shi ZL. Origin and evolution of pathogenic coronaviruses. Nature Reviews Microbiology 2019; 17 : 181–192.

3. Ramajayam R, Tan KP & Liang PH. Recent development of 3C and 3CL protease inhibitors for anti-coronavirus and anti-picornavirus drug discovery. Biochemical Society Transactions 2011; 39 : 1371–1375.

4. Wrapp D, Wang N, Corbett KS, Goldsmith JA, Hsieh CL, Abiona O et al. Cryo-EM structure of the 2019-nCoV spike in the prefusion conformation. Science 2020.

5. Wan Y, Shang J, Sun S, Tai W, Chen J, Geng Q et al. Molecular Mechanism for Antibody-Dependent Enhancement of Coronavirus Entry. J Virol 2020; 94.

6. Nelson GW, Stohlman SA & Tahara SM. High affinity interaction between nucleocapsid protein and leader/intergenic sequence of mouse hepatitis virus RNA. Journal of General Virology 2000; 81 : 181–188.

7. Stohlman SA, Baric RS, Nelson GN, Soe LH, Welter LM & Deans RJ. Specific interaction between coronavirus leader RNA and nucleocapsid protein. Journal of Virology 1988; 62 : 4288–4295.

8. Cong YY, Ulasli M, Schepers H, Mauthe M, V’kovski P, Kriegenburg F et al. Nucleocapsid Protein Recruitment to Replication-Transcription Complexes Plays a Crucial Role in Coronaviral Life Cycle. Journal of Virology 2020; 94.

9. Masters PS & Sturman LS. Background paper: Functions of the coronavirus nucleocapsid protein. Advances in Experimental Medicine and Biology 1990; 276 : 235–238.

10. McBride R, van Zyl M & Fielding BC. The Coronavirus Nucleocapsid Is a Multifunctional Protein. Viruses-Basel 2014; 6 : 2991–3018.

11. Tang TK, Wu MPJ, Chen ST, Hou MH, Hong MH, Pan FM et al. Biochemical and immunological studies of nucleocapsid proteins of severe acute respiratory syndrome and 229E human coronaviruses. Proteomics 2005; 5 : 925–937.

12. Du L, Zhao G, Lin Y, Chan C, He Y, Jiang S et al. Priming with rAAV encoding RBD of SARS-CoV S protein and boosting with RBD-specific peptides for T cell epitopes elevated humoral and cellular immune responses against SARS-CoV infection. Vaccine 2008; 26 : 1644–1651.

13. Surjit M, Liu B, Chow VTK & Lal SK. The nucleocapsid protein of severe acute respiratory syndrome-coronavirus inhibits the activity of cyclin-cyclin-dependent kinase complex and blocks S phase progression in mammalian cells. Journal of Biological Chemistry 2006; 281 : 10669–10681.

14. Hsieh PK, Chang SC, Huang CC, Lee TT, Hsiao CW, Kou YH et al. Assembly of severe acute respiratory syndrome coronavirus RNA packaging signal into virus-like particles is nucleocapsid dependent. Journal of Virology 2005; 79 : 13848–13855.

15. Ahmed SF, Quadeer AA & McKay MR. Preliminary Identification of Potential Vaccine Targets for the COVID-19 Coronavirus (SARS-CoV-2) Based on SARS-CoV Immunological Studies. Viruses 2020; 12.

16. Liu SJ, Leng CH, Lien SP, Chi HY, Huang CY, Lin CL et al. Immunological characterizations of the nucleocapsid protein based SARS vaccine candidates. Vaccine 2006; 24 : 3100–3108.

17. Shang B, Wang XY, Yuan JW, Vabret A, Wu XD, Yang RF et al. Characterization and application of monoclonal antibodies against N protein of SARS-coronavirus. Biochem Biophys Res Commun 2005; 336 : 110–117.

18. Lin Y, Shen X, Yang RF, Li YX, Ji YY, He YY et al. Identification of an epitope of SARS-coronavirus nucleocapsid protein. Cell Research 2003; 13 : 141–145.

19. Lo YS, Lin SY, Wang SM, Wang CT, Chiu YL, Huang TH et al. Oligomerization of the carboxyl terminal domain of the human coronavirus 229E nucleocapsid protein. FEBS Letters 2013; 587 : 120–127.

20. Chen IJ, Yuann JMP, Chang YM, Lin SY, Zhao J, Perlman S et al. Crystal structure-based exploration of the important role of Arg106 in the RNA-binding domain of human coronavirus OC43 nucleocapsid protein. Biochimica et Biophysica Acta - Proteins and Proteomics 2013; 1834 : 1054–1062.

21. Chang CK, Chen CMM, Chiang MH, Hsu YL & Huang TH. Transient Oligomerization of the SARS-CoV N Protein - Implication for Virus Ribonucleoprotein Packaging. PLoS ONE 2013; 8.

22. Chang CK, Sue SC, Yu TH, Hsieh CM, Tsai CK, Chiang YC et al. Modular organization of SARS coronavirus nucleocapsid protein. Journal of Biomedical Science 2006; 13 : 59–72.

23. Wootton SK, Rowland RRR & Yoo D. Phosphorylation of the porcine reproductive and respiratory syndrome virus nucleocapsid protein. Journal of Virology 2002; 76 : 10569–10576.

24. Saikatendu KS, Joseph JS, Subramanian V, Neuman BW, Buchmeier MJ, Stevens RC et al. Ribonucleocapsid formation of severe acute respiratory syndrome coronavirus through molecular action of the N-terminal domain of N protein. Journal of Virology 2007; 81 : 3913–3921.

25. Jayaram H, Fan H, Bowman BR, Ooi A, Jayaram J, Collisson EW et al. X-ray structures of the N- and C-terminal domains of a coronavirus nucleocapsid protein: Implications for nucleocapsid formation. Journal of Virology 2006; 80 : 6612–6620.

26. Fan H, Ooi A, Tan YW, Wang S, Fang S, Liu DX et al. The nucleocapsid protein of coronavirus infectious bronchitis virus: Crystal structure of its N-terminal domain and multimerization properties. Structure 2005; 13 : 1859–1868.

27. Grossoehme NE, Li L, Keane SC, Liu P, Dann Iii CE, Leibowitz JL et al. Coronavirus N Protein N-Terminal Domain (NTD) Specifically Binds the Transcriptional Regulatory Sequence (TRS) and Melts TRS-cTRS RNA Duplexes. Journal of Molecular Biology 2009; 394 : 544–557.

28. Keane SC, Lius P, Leibowitzs JL & Giedroc DP. Functional Transcriptional Regulatory Sequence (TRS) RNA binding and helix destabilizing determinants of Murine Hepatitis Virus (MHV) Nucleocapsid (N) protein. Journal of Biological Chemistry 2012; 287 : 7063–7073.

29. Tan YW, Fang S, Fan H, Lescar J & Liu DX. Amino acid residues critical for RNA-binding in the N-terminal domain of the nucleocapsid protein are essential determinants for the infectivity of coronavirus in cultured cells. Nucleic Acids Research 2006; 34 : 4816–4825.

30. Lin SM, Lin SC, Hsu JN, Chang CK, Chien CM, Wang YS et al. Structure-based stabilization of non-native protein-protein interactions of coronavirus nucleocapsid proteins in antiviral drug design. J Med Chem 2020.

31. Liebschner D, Afonine PV, Baker ML, Bunkoczi G, Chen VB, Croll TI et al. Macromolecular structure determination using X-rays, neutrons and electrons: recent developments in Phenix. Acta Crystallographica Section D-Structural Biology 2019; 75 : 861–877.

32. Wu F, Zhao S, Yu B, Chen YM, Wang W, Song ZG et al. A new coronavirus associated with human respiratory disease in China. Nature 2020.

33. de Wit E, van Doremalen N, Falzarano D & Munster VJ. SARS and MERS: recent insights into emerging coronaviruses. Nat Rev Microbiol 2016; 14 : 523–534.

34. Gorbalenya AE, Enjuanes L, Ziebuhr J & Snijder EJ. Nidovirales: Evolving the largest RNA virus genome. Virus Research 2006; 117 : 17–37.

35. Yu LL & Malik Peiris JS. Pathogenesis of severe acute respiratory syndrome. Current Opinion in Immunology 2005; 17 : 404–410.

36. Tan YJ, Lim SG & Hong W. Characterization of viral proteins encoded by the SARS-coronavirus genome. Antiviral Research 2005; 65 : 69–78.

37. Hatcher EL, Zhdanov SA, Bao Y, Blinkova O, Nawrocki EP, Ostapchuck Y et al. Virus Variation Resource - improved response to emergent viral outbreaks. Nucleic Acids Res 2017; 45 : D482–D490.

38. Robert X & Gouet P. Deciphering key features in protein structures with the new ENDscript server. Nucleic Acids Res 2014; 42 : W320–324.

39. Sievers F, Wilm A, Dineen D, Gibson TJ, Karplus K, Li WZ et al. Fast, scalable generation of high-quality protein multiple sequence alignments using Clustal Omega. Molecular Systems Biology 2011; 7.

40. Lin SY, Liu CL, Chang YM, Zhao JC, Perlman S & Hou MH. Structural Basis for the Identification of the N-Terminal Domain of Coronavirus Nucleocapsid Protein as an Antiviral Target. Journal of Medicinal Chemistry 2014; 57 : 2247–2257.

41. Zhu N, Zhang D, Wang W, Li X, Yang B, Song J et al. A Novel Coronavirus from Patients with Pneumonia in China, 2019. N Engl J Med 2020.

42. Zhou P, Yang XL, Wang XG, Hu B, Zhang L, Zhang W et al. A pneumonia outbreak associated with a new coronavirus of probable bat origin. Nature 2020.

